# Lipid-protein interactions are unique fingerprints for membrane proteins

**DOI:** 10.1101/191486

**Authors:** Valentina Corradi, Eduardo Mendez-Villuendas, Helgi I. Ingólfsson, Ruo-Xu Gu, Iwona Siuda, Manuel N. Melo, Anastassiia Moussatova, Christine Degagné, Besian I. Sejdiu, Gurpreet Singh, Tsjerk A. Wassenaar, Karelia Delgado Magnero, Siewert J. Marrink, D. Peter Tieleman

## Abstract

Cell membranes contain hundreds of different proteins and lipids in an asymmetric arrangement. Understanding the lateral organization principles of these complex mixtures is essential for life and health. However, our current understanding of the detailed organization of cell membranes remains rather elusive, owing to the lack of experimental methods suitable for studying these fluctuating nanoscale assemblies of lipids and proteins with the required spatiotemporal resolution. Here, we use molecular dynamics simulations to characterize the lipid environment of ten membrane proteins. To provide a realistic lipid environment, the proteins are embedded in a model plasma membrane, where more than 60 lipid species are represented, asymmetrically distributed between leaflets. The simulations detail how each protein modulates its local lipid environment through local lipid composition, thickness, curvature and lipid dynamics. Our results provide a molecular glimpse of the complexity of lipid-protein interactions, with potentially far reaching implications for the overall organization of the cell membrane.

## Introduction

Lipids and proteins are the major components of all biological membranes, which play crucial roles with respect to the structure and function of the cell. The hydrophilic headgroup and the hydrophobic acyl tails of lipids allow their assembly into lamellar structures, thus separating the interior from the exterior of the cell, as in the case of the plasma membrane, or segregating different intracellular compartments from the cytosol. Membrane proteins carry out a large variety of functions. Integral membrane proteins can act as receptors, involved in signal transduction, or as channels or transporters, thus involved in the transfer of solutes from one side of the membrane to the other. Membrane proteins can also promote the interaction between cells, intracellular compartments or large macromolecular complexes, or can act as enzymes.

A complex lipid-protein interplay takes place in the membrane.^1-2^ Lipids do not simply provide the matrix were proteins are embedded but can actively participate to the regulation of protein activity, trafficking and localization.^2^ Proteins, on the other hand, do shape lipids by inducing membrane deformations and lipid sorting mechanisms.^3-4^ The complexity of such interplay is also a consequence of the large variety of lipid types and their asymmetric distribution found in biological membranes.^2^ Thus, lipid-protein interplay occurs via multiple mechanisms, which include interactions that can be (i) specific, where a clear binding site for a given lipid or headgroup can be identified, or (ii) non-specific, where lipids act as a medium and physical properties like thickness, fluidity, or curvature regulate protein function.^4-5^

The characterization of lipid-protein interactions provides crucial details for a better understanding of the biological activity of a given membrane protein. In the last few decades, several experimental and computational techniques have been used to answer questions related to the identification of lipid binding sites on the protein surface, the type of lipids found associated with the protein, and how such lipids influence protein function.^6^ X-ray crystallography and electron crystallization have identified a number of lipids strongly bound to proteins as these lipids have to survive the crystallization process.^7-10^ Lipid binding to membrane proteins and the local lipid composition in proximity of the protein can also be studied using fluorescence methods.^11-13^ Biophysical studies of lipid-protein interactions have also used nanodiscs, discoidal membranes with a diameter of 8-17 nm enclosed by helical scaffolding proteins, which allows controlling the lipid composition.^14-15^ Recently, the application of detergent-free approaches that use specific copolymers to extract proteins and native lipids into nanodiscs has provided a new tool to characterize the lipid environment of a given membrane protein.^16^ Quantitative analysis and identification of native lipid species tightly associated with membrane proteins can be achieved via mass-spectroscopy, where lipid-protein complexes can be solubilized in non-ionic detergents to provide resistance to the electron spray ionization step, thus allowing for stable complexes in gas phase.^17^ These techniques focus on strong interactions and although some are qualitative, they do not give high spatial resolution.

Computer simulations have also been extensively used to study membranes and membrane proteins systems at an atomistic or near atomistic level of detail.^18-22^ Such simulations describe in detail the motion of lipids, proteins and other particles, as well as their interactions, providing information on the structure, dynamics and thermodynamics of the system. Molecular dynamics (MD) simulations that use coarse-grained (CG) models such as the Martini force field^23^ are now routinely applied not only to study physical properties of lipid bilayers, but also to investigate specific and non-specific lipid-protein interactions (reviewed in ^18, 23-25^). Membrane proteins are now simulated not only in controlled lipid environments that match the lipid composition used in experiments, but also in more realistic bilayers that mimic the natural lipid environment.^25-34^

In this paper we use CG MD to study the lipid distribution and membrane properties of a complex plasma lipid mixture^35^ and representatives of ten diverse plasma membrane protein families (). The proteins considered are: aquaporin 1 (AQP1), prostaglandin H2 synthase (COX1), dopamine transporter (DAT), epidermal growth factor (EGFR), AMPA-sensitive glutamate receptor (GluA2), glucose transporter (GLUT1), voltage-dependent Shaker potassium channel 1.2 (Kv1.2), sodium, potassium pump (Na,K-ATPase), δ-opioid receptor (δ-OPR), and P-glycoprotein (P-gp). These proteins include transporters, channels, enzymes, and receptors, and represent different quaternary structures and sizes, as well as monotopic membrane proteins. As shown in **Error! Reference source not found.**, the actual simulation system contains four copies of the same protein, to increase statistics in a computationally efficient way and to have an additional estimate of statistical errors independent of time correlations. **Error! Reference source not found.** also shows the general set up with a complex asymmetric lipid mixture consisting of more than 60 lipid types.^35^

Based on 30 μs simulations for each system (Figure 2 - Figure Supplements 1-3), we describe the distinctive nature of the lipid environment surrounding each protein, analyzing lipid distribution, cholesterol dynamics, and membrane properties including thickness and curvature. The results show a rich variety of lipid-protein interactions and protein effects on membrane physics, emphasizing the importance of not just tightly-bound lipids but the overall structure of the lipidprotein matrix.

**Figure 1.**
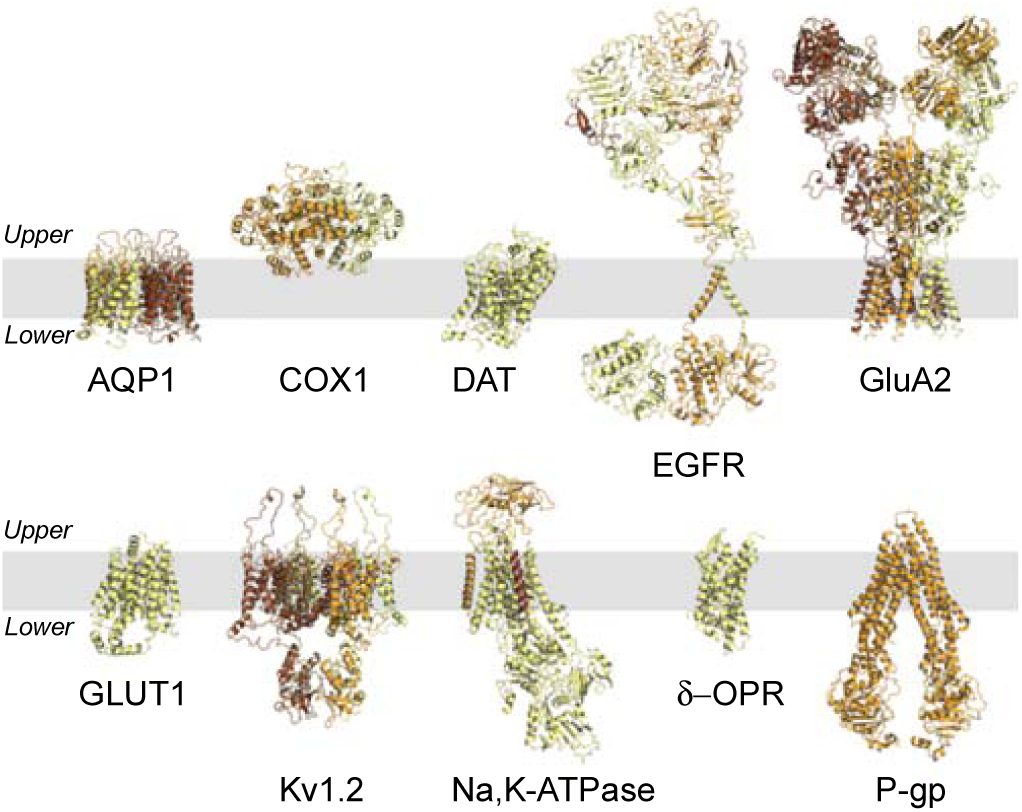
Membrane Proteins. Atomistic structures of the membrane proteins selected for this study. Each protein is shown as cartoons, coloured from light yellow to brown when multiple chains are present. The gray-shaded area represents the hydrophobic region of the membrane.

**Figure 2.**
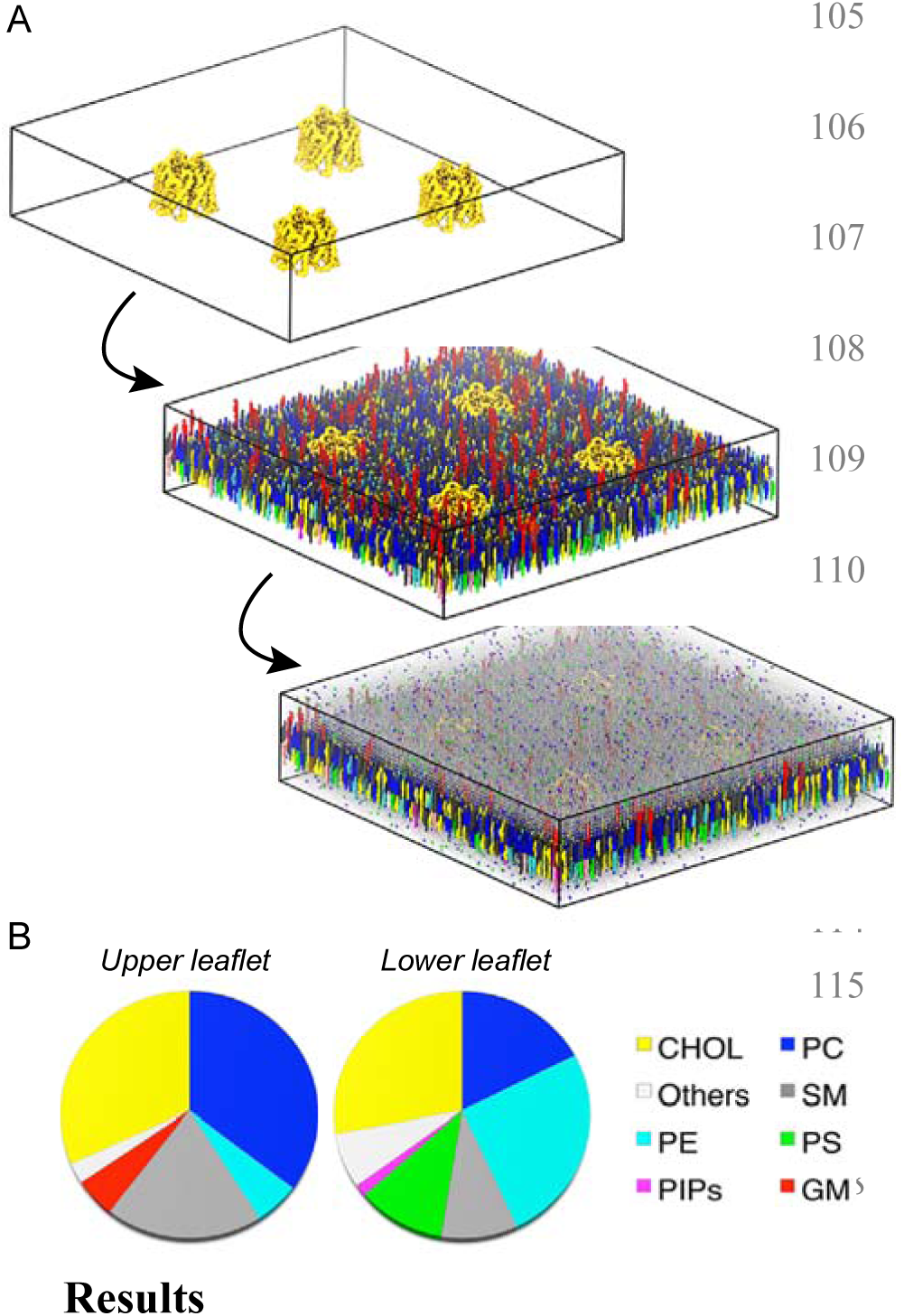
System setup and lipid composition. (A) For a given system, four protein molecules are placed in a simulation box of ca. 42 x 42 nm in x and y. Lipids, water and ions are added using *insane.*^36^ For clarity, the setup is shown as a two-steps process. (B) Lipid composition of the main headgroup types in upper and lower leaflet, color-coded as in A.

### Annular lipid shells have unique lipid compositions

To analyze the lipid surroundings of each protein, we calculated 2D lateral density maps (see Methods), averaged over the four copies of the protein present in the simulation cell and considering the last 5 μs of the trajectory. Given the complex composition of our membrane, we grouped the lipids into four major classes, i.e poly-unsaturated (PU) lipids, fully-saturated (FS) lipids, cholesterol (CHOL), and all remaining lipids (Others). The PU, FS, and CHOL density maps are shown in Figure 3 for all the proteins, averaged over the four protein molecules of each system. The full set of data for all systems is shown in Figure 3-Figure Supplement 1 (PU, FS, CHOL groups) and 2 (Others group). We observe a rich spectrum of possible modes of interactions. These include non-specific binding, as shown, for example, by the broad distribution of PU and FS lipids found, in both leaflets, near many proteins, including AQP1, DAT, EGFR, GluA2, GLUT1, Na,K-ATPase and δ-OPR. These features are somewhat leaflet-specific as they depend on protein structure and lipid composition, both of which are asymmetric. In the upper leaflet, for example, we notice regions of strong FS enrichment in contact with the proteins. Such regions are usually more localized than PU enriched regions and are often coupled with smaller, yet still highly localized, FS enriched regions in the lower leaflet. This behaviour can be seen for AQP1, DAT, EGFR, δ-OPR, and even for the monotopic COX1, which is only partially embedded in the upper leaflet. The size and shape of FS lipid regions differ from protein to protein, and in many cases create a discontinuous ring around the transmembrane domains, as in the case of AQP1, GluA2, and Kv1.2, which are homotetramers. In the lower leaflet, PU enrichment is often observed near the proteins, and particularly noticeable around, for example, DAT, GluA2, GLUT1, and Na,K-ATPase. Some proteins clearly induce a sharp partitioning of the different lipid classes. This is the case for GluA2 and PU lipids in the lower leaflet, or P-gp, where in the upper leaflet we observe a clear distinction between the side of the transmembrane domains in contact with PU lipids and the side in contact with FS lipids, while in the lower leaflet there is an obvious preference for PU lipids. Kv1.2 is another striking example of how the same lipid class (PU) can be asymmetrically distributed between leaflets, and symmetrically distributed around the protein within the same leaflet, due to the homo-tetrameric nature of the channel and possibly linked to a more specific type of binding.

**Figure 3.**
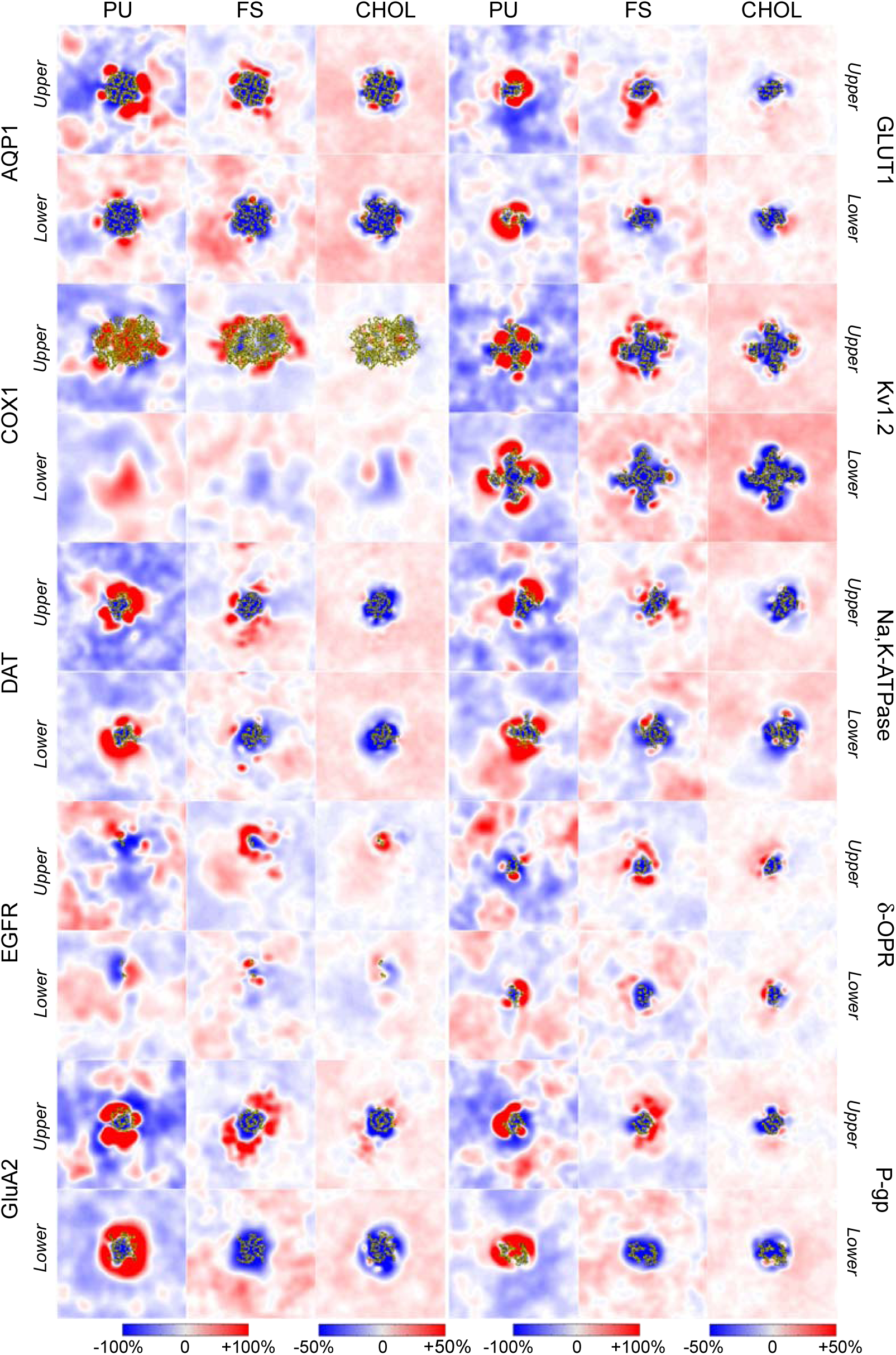
Lipid density distribution. Lipid density analysis of the poly-unsaturated (PU), fully-saturated (FS), and cholesterol (CHOL) classes. The lipid density is represented by x and y 2D maps, averaged between 25 to 30 μs and over the four protein molecules of a given system. The maps are colored by relative enrichment (red) or depletion (blue), calculated with respect to the average (white) density of a given class. The portion of the protein intersecting the upper and lower surfaces used for the calculation is shown in yellow ribbons.

**Figure 4.**
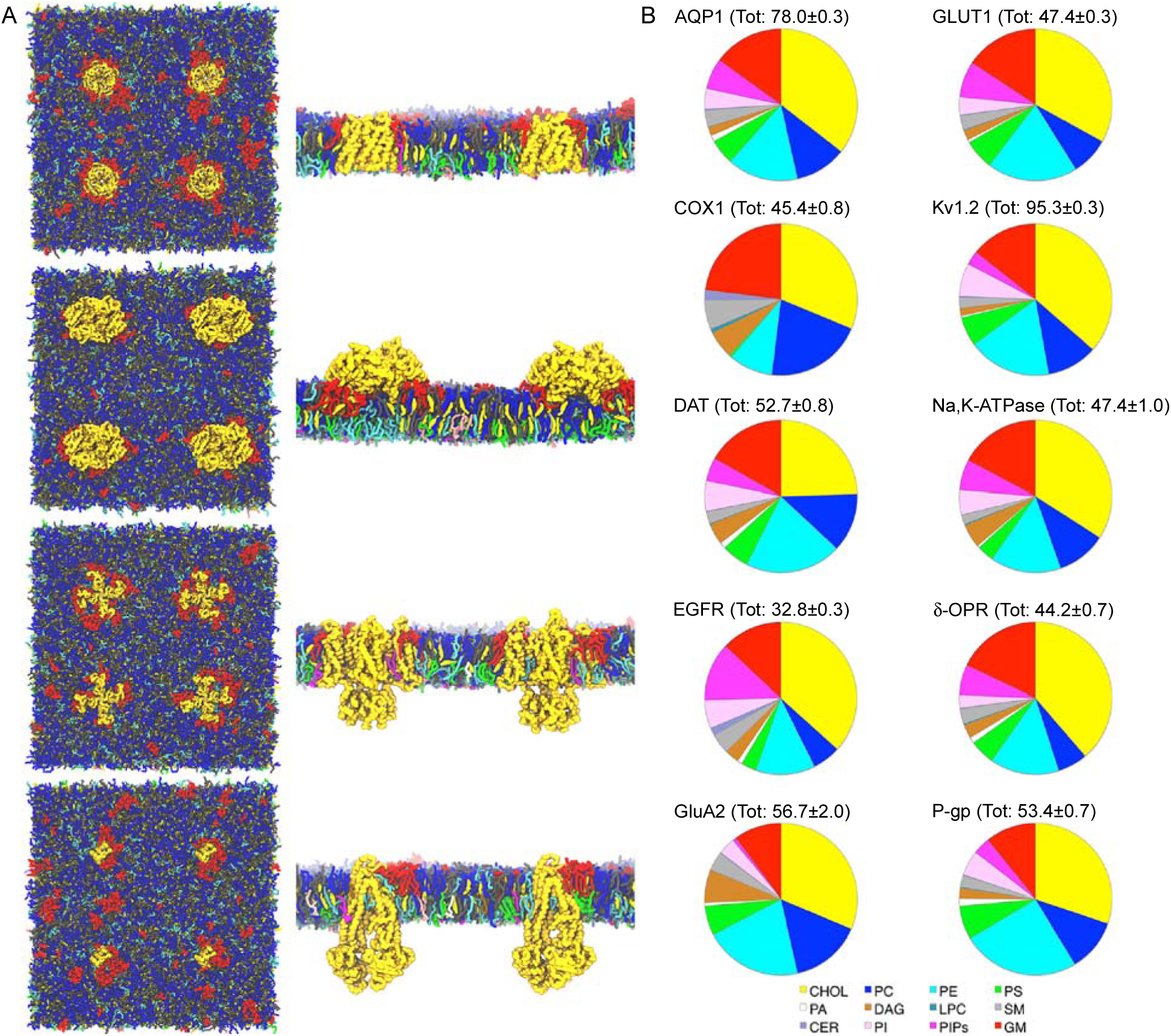
First lipid shell composition. (A) Snapshots of the upper leaflet of the AQP1, COX1, Kv1.2 and P-gp system and the side view (with lipids clipped for clarity) of two of the four protein molecules to show the lipids arrangement around the proteins transmembrane domains. (B) Pie charts showing the lipid headgroup composition of the first lipid shell for the ten system. The total number of lipids found within the selected 0.7 nm cutoff is reported in parenthesis as average number of lipids obtained from the four protein copies of each system (the standard error is reported).

Monotopic proteins are also capable of inducing a clear separation in the distribution of lipid classes, even in the leaflet they are not directly bound to. For COX1, for example, in the lower leaflet the enrichment of PU lipids stands against the depletion of FS lipids and CHOL underneath the protein, partially embedded only in the upper leaflet. In addition to a non-specific, broad distribution of lipid classes, we detect example of specific binding, especially for the CHOL class. CHOL is the most represented component of the plasma membrane mixture, thus associated with a more even distribution compared to the PU and FS classes. However, the corresponding CHOL-2D density maps reveal site of specific cholesterol binding. AQP1, for example, clearly shows very specific binding of cholesterol at the interface between monomers, but an indication of specific cholesterol binding can be detected in DAT, GLUT1, Kv1.2, Na,K-ATPase and δ-OPR as well. Overall, many of the features described above, which are obtained from 5 µs me windows averaging, can be seen at different timescales, as shown in the movie files representing the lipid distribution of the AQP1 system at different averaging times (200 ns and 2000 ns (movies S1-S2).

According to the analysis discussed above, each protein is associated with a unique lipid distribution. However, common features can be detected, as, for example, regions near the proteins enriched in FS lipids in the upper leaflet, the accumulation of PU lipids around most proteins, or confined regions of cholesterol binding. We further investigated the presence, across the systems, of patterns in lipid distribution by (i) considering the lipid composition of the first lipid shell surrounding the protein (within a 0.7 nm cutoff), and (ii) by quantifying the enrichment (or depletion) in a given lipid class in the immediate proximity of the proteins using the depletion-enrichment (D-E) index analysis.

As the size of the protein transmembrane domains is different from protein to protein, the total number of lipids found within the first lipid shell varies from ca. 32 as in EGFR (whose transmembrane domain consists of only two helices) to ca. 78 for AQP1 and ca. 95 for Kv1.2 (which are tetrameric proteins) (**Error! Reference source not found.**).

Common to all the lipid shells (averaged over the four protein molecules of a given system) here analyzed is the prevalence of CHOL as well as the presence of GM lipids (with COX1 showing the highest GM fraction). The fraction of PC and PE lipids (the most abundant phospholipid in the upper and lower leaflet, respectively) changes considerably from protein to protein, with GluA2 and P-gp showing higher content of PE lipids. From the lower leaflet, PIP, PI and PS lipids contribute to the composition of the lipid shell in a similar way for many proteins, with the exception of EGFR (where the PIP lipids content is higher than the PI content), COX1 (which is only partially embedded in the upper leaflet) and GluA2 (for which the fraction of PIP lipids is significantly smaller than other proteins). DAG and SM lipids can also contribute to smaller fractions of the lipid shell.

We then quantified the depletion or enrichment of a given lipid type within this first shell and up to 2.1 nm from the proteins. According to our definition of the D-E index, values larger than 1 indicate enrichment of a given lipid group within a given distance cut-off, while values smaller than 1 indicate depletion. Table 1 summarizes the results for the four lipid classes also used in the lipid density analysis (i.e. PU, FS, CHOL and Others). A clear enrichment of PU lipids is present within the shortest distance (with the exception of EGFR), as well as FS lipids in the case of several systems (Table 1 and Supplementary File 2, Tables 1 - Table Supplements 1 and 2). This revealed additional patterns of lipid organization. For example, the enrichment in FS lipids is linked to the FS-GM lipids enrichment in the upper leaflet (Supplementary File 2, Table 1 Table Supplements 1 and 2). Together with the enrichment of GM lipids, we observe a depletion of PC lipids, despite being the most abundant phospholipid type in the upper leaflet. In the lower leaflet, enrichment of PIP lipids is common to all the simulation systems (with the exception of COX1). Interestingly, the enrichment of PIP lipids is also accompanied by a certain degree of enrichment of other negatively charged lipids, like PI lipids (Supplementary File 2, Table 1 Tables Supplements 1 and 2).

**Table 1.** Depletion-Enrichment (D-E) values for Fully-Saturated (FS), Poly-Unsaturated (PU), Cholesterol (CHOL) and Others (neither PU nor FS or CHOL) lipid classes.

|  | FS |  |  | PU |  |  | CHOL |  |  | Others |  |  |
| --- | --- | --- | --- | --- | --- | --- | --- | --- | --- | --- | --- | --- |
|  | Distance cut-off (nm) |  |  | Distance cut-off (nm) |  |  | Distance cut-off (nm) |  |  | Distance cut-off (nm) |  |  |
|  | 0.7 | 1.4 | 2.1 | 0.7 | 1.4 | 2.1 | 0.7 | 1.4 | 2.1 | 0.7 | 1.4 | 2.1 |
| AQP1 | 1.26±0.16 | 1.26±0.10 | 1.18±0.08 | 1.76±0.60 | 1.53±0.49 | 1.37±0.41 | 1.18±0.04 | 1.07±0.04 | 1.03±0.04 | 0.82±0.02 | 0.89±0.01 | 0.93±0.01 |
| COX1 | 1.70±0.29 | 1.76±0.24 | 1.74±0.21 | 1.33±0.76 | 0.96±0.41 | 0.74±0.25 | 1.04±0.02 | 1.06±0.03 | 1.06±0.01 | 0.84±0.07 | 0.84±0.05 | 0.86±0.04 |
| DAT | 1.26±0.22 | 1.24±0.14 | 1.18±0.10 | 4.16±1.34 | 3.01±0.92 | 2.34±0.66 | 0.81±0.07 | 0.88±0.04 | 0.92±0.03 | 0.87±0.02 | 0.91±0.02 | 0.94±0.01 |
| EGFR | 1.11±0.41 | 1.20±0.09 | 1.15±0.05 | 0.89±0.14 | 1.20±0.07 | 1.17±0.09 | 1.22±0.02 | 0.94±0.01 | 0.95±0.01 | 0.87±0.07 | 0.98±0.01 | 0.99±0.01 |
| GluA2 | 0.68±0.25 | 1.18±0.12 | 1.23±0.11 | 4.57±1.31 | 3.13±0.72 | 2.31±0.49 | 1.04±0.11 | 0.98±0.06 | 0.98±0.04 | 0.83±0.03 | 0.86±0.03 | 0.90±0.04 |
| GluT1 | 1.05±0.27 | 1.21±0.21 | 1.18±0.15 | 3.89±1.17 | 2.74±0.76 | 2.08±0.54 | 1.10±0.07 | 1.01±0.04 | 1.00±0.03 | 0.77±0.05 | 0.86±0.03 | 0.91±0.01 |
| Kv1.2 | 0.90±0.15 | 1.17±0.14 | 1.13±0.09 | 2.96±0.30 | 2.51±0.32 | 2.06±0.26 | 1.21±0.04 | 1.03±0.02 | 1.00±0.01 | 0.79±0.04 | 0.87±0.04 | 0.92±0.03 |
| Na,K-ATPase | 1.40±0.15 | 1.21±0.07 | 1.10±0.06 | 2.99±1.09 | 2.48±0.76 | 2.01±0.50 | 1.13±0.03 | 0.98±0.02 | 0.97±0.03 | 0.75±0.04 | 0.89±0.03 | 0.94±0.03 |
| δ-OPR | 0.99±0.24 | 1.27±0.16 | 1.23±0.14 | 2.10±0.76 | 1.51±0.44 | 1.25±0.32 | 1.29±0.01 | 1.12±0.02 | 1.08±0.01 | 0.79±0.02 | 0.86±0.04 | 0.90±0.04 |
| P-gp | 0.87±0.19 | 1.04±0.21 | 1.09±0.12 | 4.79±0.80 | 3.22±0.54 | 2.33±0.39 | 1.00±0.01 | 0.98±0.02 | 0.98±0.02 | 0.80±0.04 | 0.88±0.03 | 0.92±0.02 |

For each lipid class, the values obtained for three distance cut-offs (0.7, 1.4, and 2.1 nm) from the proteins are shown. The values are the average obtained from the four protein molecules of each system.

### Membrane thickness deformations are highly non-uniform

Membrane proteins are known to perturb the thickness of the membrane to optimize their embedding.^37^ There are several possible mechanisms for membrane thickness to vary. Variations in thickness have been observed even in pure bilayers in some of the earliest membrane protein simulations,^38^ but in complex mixtures an obvious mechanism is a non-uniform distribution of lipids. The lipid enrichment and depletion patterns revealed in the previous section are expected to impact the local membrane thickness. **Error! Reference source not found.** shows the 2D thickness distribution of our plasma membrane model, divided in upper (outer) and lower (inner) leaflet as well as the total thickness for four selected proteins, i.e. AQP1 (as an example of a membrane-spanning homotetramer with a cylindrical-like structure); COX1 (chosen as an example of monotopic membrane protein); Kv1.2 (as an example of a homotetramer with a more complex structure than AQP1); P-gp (as an example of proteins for which lipids also act as substrates).^39^

For these four selected proteins the 2D thickness maps of Figure 5 are shown as the average among the four protein molecules in each system. Additional data on each protein copy for all the systems is given in Figure 5 - Figure Supplement 1.

**Figure 5.**
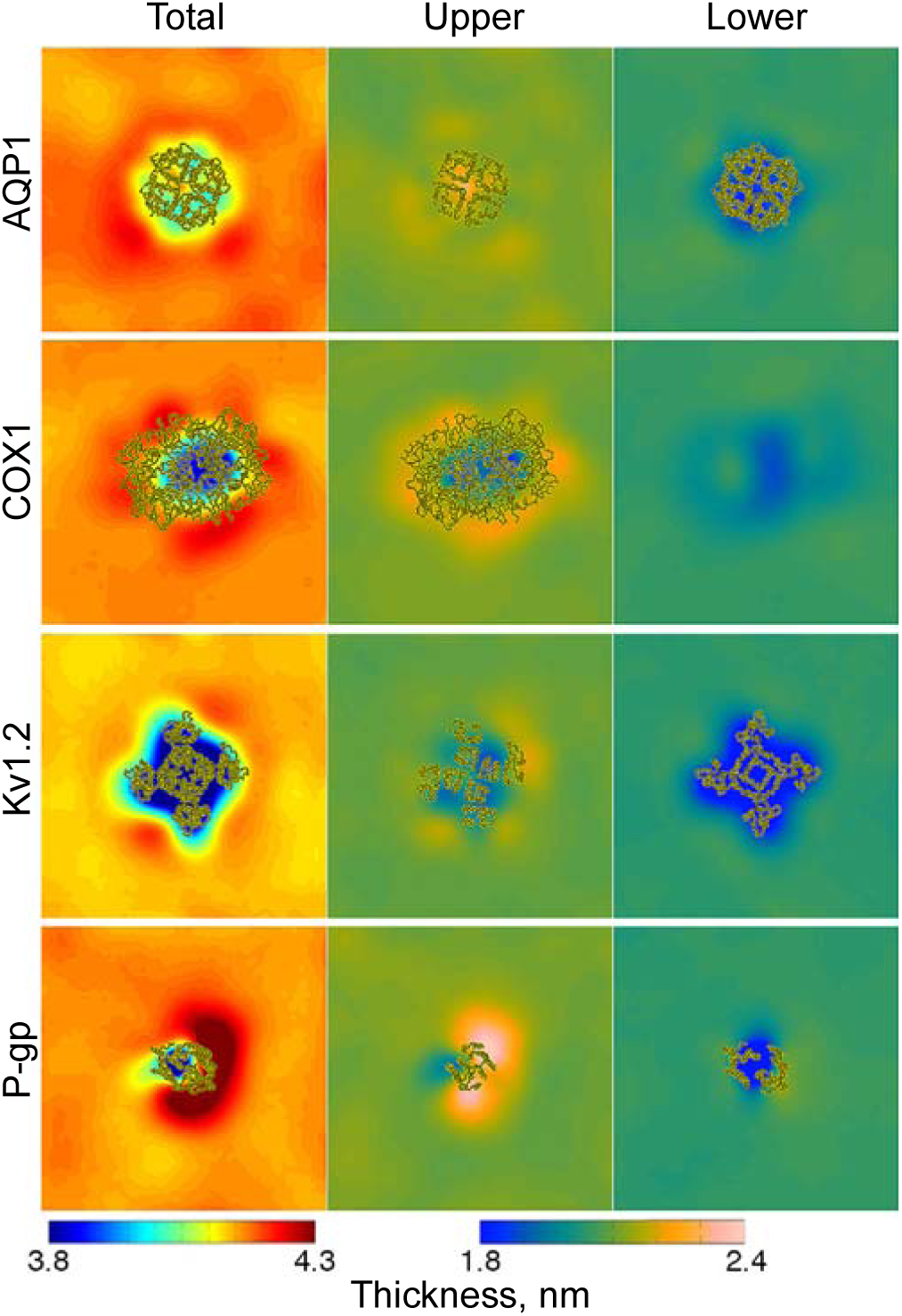
Membrane thickness. Membrane thickness. For four selected systems (AQP1, COX1, Kv1.2, and P-gp) membrane thickness is shown as x and y 2D maps, averaged between 25 to 30 μs and over the four protein molecules of a given system. Total thickness, i.e. the distance calculated between the upper and lower surfaces used for the analyses, is shown color-coded according to a 3.8 to 4.3 nm range. Thickness maps for the upper leaflet and lower leaflet are shown on a different color scale, ranging from 1.8 to 2.4 nm. The portion of the protein intersecting the different surfaces is shown in yellow ribbons.

Overall, the extent of spatial perturbation is quite significant, as essentially the entire system is affected by the presence of the proteins. As expected, the strongest effects are near a protein molecule, and depend strongly on the specific nature of the protein, as well as on its interactions with local lipids. Thinning of the membrane near the proteins, linked to their hydrophobic belt, can be uniformly distributed around the transmembrane region of structurally symmetric proteins, as for the tetrameric AQP1 and Kv1.2 (**Error! Reference source not found.**). P-gp is an example of an asymmetric thickness profile, where higher thickness is detected only on one side of its transmembrane domains (**Error! Reference source not found.**). Depending on the shape of the protein, the effects on the upper and lower leaflet of the membrane are different, with COX1 as an extreme case as it is bound only to one side of the bilayer. Yet despite this, the opposite leaflet still couples with changes in thickness of the binding leaflet. In general, the thickness is higher for the upper leaflet than for the lower one, a result of the asymmetric lipid composition, with the lower leaflet enriched in unsaturated lipids, and the upper leaflet enriched in more saturated and longer tailed lipids (as per bilayer composition).^35^ However, strong variations in the overall geometry of the bilayer are observed as a function of distance from the proteins, with deformations spanning the first few nm from the proteins, and possibly extending to reach equivalent neighbouring protein molecules. The total thickness maps reveal a thinning of the membrane near many proteins, together with highly confined region of increased thickness, as a reminder that the shape and size of the protein hydrophobic belt might not be uniform around the transmembrane domains, thus affecting the local lipid distribution.^40^ As shown for the AQP1 case (see movie S3, obtained by averaging over 200 ns time windows), these features persist at different timescales.

### Membrane curvature effects are long ranged

Proteins remodel membranes by inducing changes in thickness and lipid composition, reflected in changes of the local membrane curvature. Here, we describe the overall bending of the membrane by means of Mean and Gaussian curvatures (K_M_ and K_G_, respectively), calculated for the upper, middle and lower surfaces, and shown with respect to the normal defined by the upper plane (see Methods). While K_M_ is an indication of the extent of bending, K_G_ provides information on the membrane topology, being dependent on the sign of the principal curvatures.

For a given surface, positive K_M_ values indicate a convex surface, while negative values or zero indicate a concave and a flat surface, respectively. K_G_ negative values are associated with surface saddles, while positive values or zero values correspond to a spherical and a cylindrical topology, respectively. Figure 6 shows the results for the four selected systems, while Figure 6 Figures Supplements 1 and 2 provide K_M_ and K_G_ details on all the systems with the four proteins. Each simulation system, regardless of symmetry and size of the transmembrane domains, clearly reveals a complex curvature landscape, with a strong coupling between surfaces even for proteins only partially embedded in the membrane, as COX1 (**Error! Reference source not found.**).

**Figure 6.**
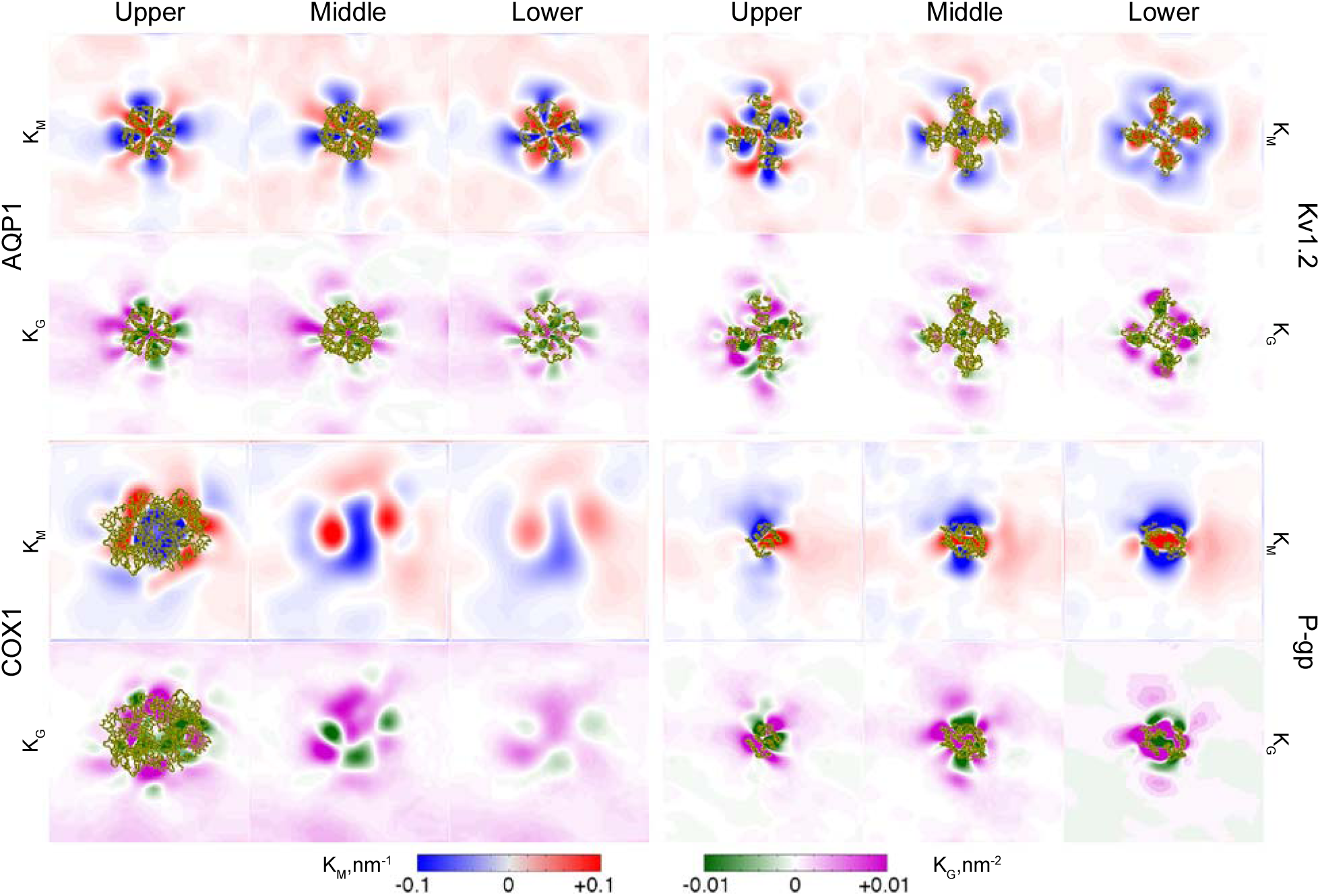
Membrane curvature. 2D maps of the Mean and Gaussian curvature (K_M_ and K_G_, respectively) of the four selected systems (AQP1, COX1, Kv1.2, and P-gp), averaged between 25 to 30 μs and over the four protein molecules of a given system. The upper, middle and lower surfaces used to calculate thickness were employed to derive the values of K_M_ and K_G_, defined with respect to the normal of the upper surface. The portion of the protein intersecting the three different surfaces used for the calculation is shown in yellow ribbons.

Overall, K_G_ is, on average, shifted towards positive or zero values, while K_M_ shows wide regions of positive and negative curvature, with the stronger changes located in close proximity of the proteins (**Error! Reference source not found.** and Figure 6 Figure Supplement 2), where features of the local curvature are maintained for the full length of the simulation (as shown for APQ1 in SI, movie S4). In addition, among the four proteins in a given system, the curvature profile is, qualitatively, very similar (Figure 6 Figures Supplements 1 and 2). This is particularly noticeable for larger and symmetric transmembrane domains, e.g. AQP1 and Kv1.2, as well as for proteins not characterized by such a high degree of structural symmetry, like P-gp (Figure 6 and Figure 6 Figures Supplements 1 and 2). To connect the observed curvatures more directly to the structure of the proteins, we identified sample structural features of two proteins (AQP1 and P-gp) that correlate with the profile of the K_M_ curvature map (**Error! Reference source not found.**). The different structures of the two proteins cause very different effects in the membrane: In the case of AQP1, the characteristic pattern of positive and negative curvature of the upper (and middle) surface appears linked to the interface regions between monomers and to the presence of long extracellular loops, while for P-gp the shorter extracellular ends of TM3 and TM4 together with the presence of positively charged residues (Arg206 and Lys209) relate to the nearby negative curvature.

### Rate of cholesterol flip-flop strongly depends on protein-lipid environment

Cholesterol is the most abundant component of eukaryotic plasma membranes. Its ability of redistributing between domains of different composition in a given leaflet, and between leaflets of asymmetric composition^41-42^ is of crucial importance in regulating and controlling both protein function and membrane properties. We looked in particular at how proteins affect the distribution of cholesterol across the leaflets (flip-flop) as multiple mechanisms can be involved. Cholesterol molecules might, for example: (i) interact with a given protein site for several microseconds before slowly climbing along the protein surface, flipping to the opposite leaflet and moving further away, where flip-flop events will occur more freely (**Error! Reference source not found.**A); (ii) remain bound to a given site for longer timescales, thus refraining that cholesterol from flipflopping (**Error! Reference source not found.**B); or (iii) slowly reduce the number of flip-flop events as it approaches a protein molecule and establishes stronger interactions within a given site (**Error! Reference source not found.**C).

To better characterize the cholesterol dynamics in the presence of proteins, we defined flipflop and flip-in events and explored how they vary at different distance from the proteins’ transmembrane domains (Table 2, and Supplementary File 2, Table 2 Table Supplement 1). The rate of cholesterol flip-flop in simulations of the pure plasma mixture was estimated at ca. 6.5 x 10^6^ s^-1^.^35^ Similar values are here met at different distances from the proteins, depending on the protein in question. Using a 5 Å distance bin width, at a distance greater than 60 Å (which we define here as bulk), over a 5 μs time period, the rate of cholesterol flip-flop ranges from ca.20 (GluA2) to 33-34 (e.g. AQP1) events per molecule (Table 2), yielding rates that span from 5.2 x 10^6^ s^-1^ (GluA2) to 6.8 x 10^6^ s^-1^ (AQP1), similarly to a pure plasma membrane.

**Table 2.**
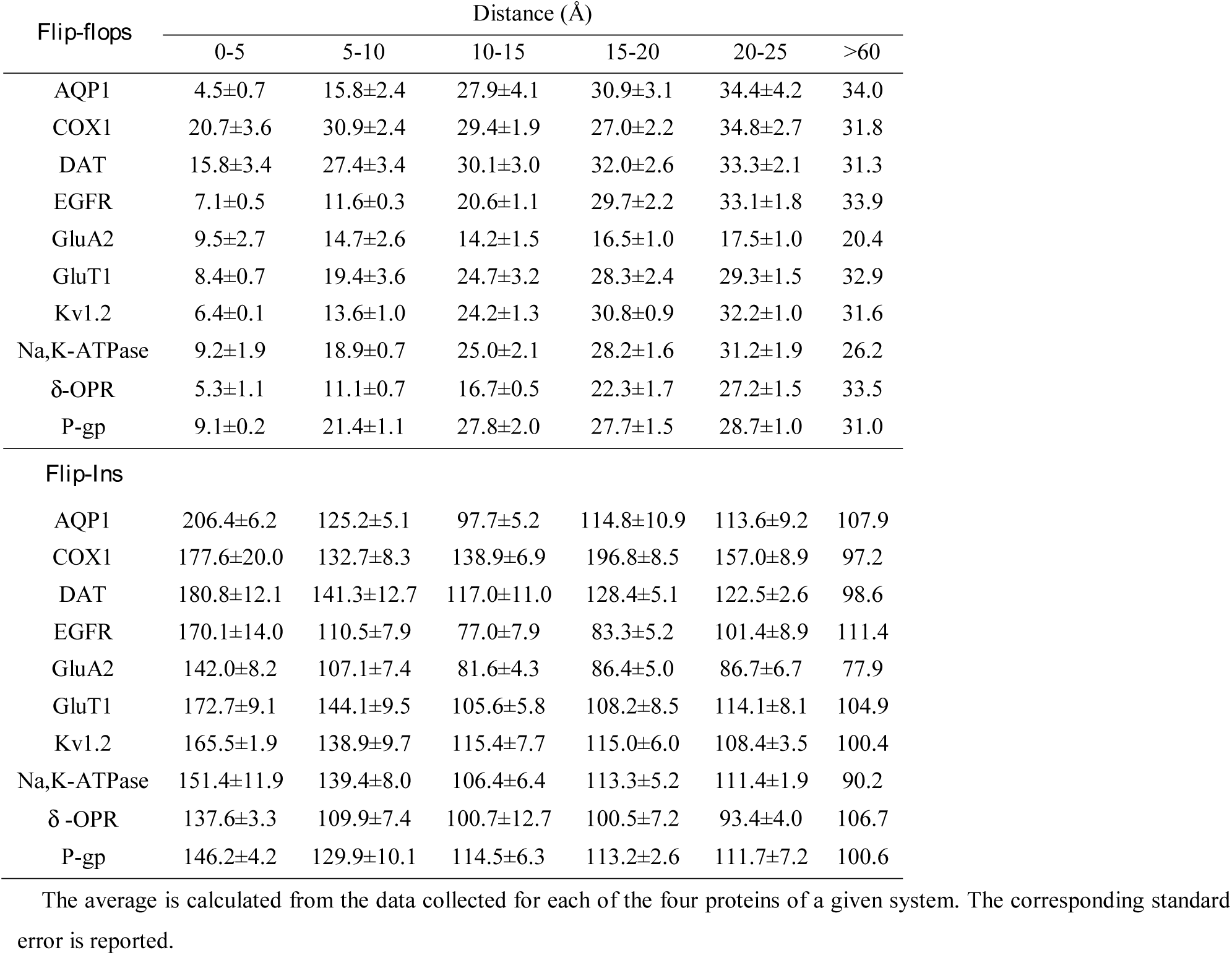
Cholesterol flip-flop and flip-in events. For all the systems, the average number of flip-flop and flip-in events per cholesterol molecule over the last 5 s of the simulation is shown as a function of distance from the proteins.

| *Flip-flops* | Distance ($\text{\AA}$) | | | | | |
| --- | --- | --- | --- | --- | --- | --- |
|  | 0-5 | 5-10 | 10-15 | 15-20 | 20-25 | >60 |
| AQP1 | 4.5±0.7 | 15.8±2.4 | 27.9±4.1 | 30.9±3.1 | 34.4±4.2 | 34.0 |
| COX1 | 20.7±3.6 | 30.9±2.4 | 29.4±1.9 | 27.0±2.2 | 34.8±2.7 | 31.8 |
| DAT | 15.8±3.4 | 27.4±3.4 | 30.1±3.0 | 32.0±2.6 | 33.3±2.1 | 31.3 |
| EGFR | 7.1±0.5 | 11.6±0.3 | 20.6±1.1 | 29.7±2.2 | 33.1±1.8 | 33.9 |
| GluA2 | 9.5±2.7 | 14.7±2.6 | 14.2±1.5 | 16.5±1.0 | 17.5±1.0 | 20.4 |
| GluT1 | 8.4±0.7 | 19.4±3.6 | 24.7±3.2 | 28.3±2.4 | 29.3±1.5 | 32.9 |
| Kv1.2 | 6.4±0.1 | 13.6±1.0 | 24.2±1.3 | 30.8±0.9 | 32.2±1.0 | 31.6 |
| Na,K-ATPase | 9.2±1.9 | 18.9±0.7 | 25.0±2.1 | 28.2±1.6 | 31.2±1.9 | 26.2 |
| δ-OPR | 5.3±1.1 | 11.1±0.7 | 16.7±0.5 | 22.3±1.7 | 27.2±1.5 | 33.5 |
| P-gp | 9.1±0.2 | 21.4±1.1 | 27.8±2.0 | 27.7±1.5 | 28.7±1.0 | 31.0 |
| *Flip-Ins* |  |  |  |  |  |  |
| AQP1 | 206.4±6.2 | 125.2±5.1 | 97.7±5.2 | 114.8±10.9 | 113.6±9.2 | 107.9 |
| COX1 | 177.6±20.0 | 132.7±8.3 | 138.9±6.9 | 196.8±8.5 | 157.0±8.9 | 97.2 |
| DAT | 180.8±12.1 | 141.3±12.7 | 117.0±11.0 | 128.4±5.1 | 122.5±2.6 | 98.6 |
| EGFR | 170.1±14.0 | 110.5±7.9 | 77.0±7.9 | 83.3±5.2 | 101.4±8.9 | 111.4 |
| GluA2 | 142.0±8.2 | 107.1±7.4 | 81.6±4.3 | 86.4±5.0 | 86.7±6.7 | 77.9 |
| GluT1 | 172.7±9.1 | 144.1±9.5 | 105.6±5.8 | 108.2±8.5 | 114.1±8.1 | 104.9 |
| Kv1.2 | 165.5±1.9 | 138.9±9.7 | 115.4±7.7 | 115.0±6.0 | 108.4±3.5 | 100.4 |
| Na,K-ATPase | 151.4±11.9 | 139.4±8.0 | 106.4±6.4 | 113.3±5.2 | 111.4±1.9 | 90.2 |
| δ-OPR | 137.6±3.3 | 109.9±7.4 | 100.7±12.7 | 100.5±7.2 | 93.4±4.0 | 106.7 |
| P-gp | 146.2±4.2 | 129.9±10.1 | 114.5±6.3 | 113.2±2.6 | 111.7±7.2 | 100.6 |

The average is calculated from the data collected for each of the four proteins of a given system. The corresponding standard error is reported.

More in detail, many systems, e.g. AQP1 and Kv1.2, show such bulk values after ca. 10 to 20 Å distance from the proteins. Others, as P-gp, affect the flip-flop rate at larger distances, and bulk values are met after ca. 25 Å, while for the membrane associated COX1, the effects on cholesterol dynamics across the leaflets are milder, and bulk values can be seen at much smaller distances (exceeding 5 Å). Overall, for all the systems, the slowest flip-flop rates are found in the immediate proximity of the proteins (0-5 Å), with significantly slower rates observed for example for AQP1 (ca. 4.5 flip-flop events per cholesterol molecule, yielding a 9 x 10^5^ s^-1^ flip-flop rate). The lower flip-flop rates for regions adjacent to the proteins are associated with considerably higher flip-in (transition from upper or lower leaflet to bilayer middle) rates (Table 2 and Supplementary File 2, Table 2 Table Supplement 1). Within the first 5 Å from the proteins, for example, the number of flip-in events for AQP1 is ca. 206.4, yielding rates of ca. 4.1 x 10^7^ s^-1^, respectively. When plotted as 2D map, flip-flop events are detected over the full length of a given simulation box, independently from the size and topology of the transmembrane domains. Smaller, localized regions of higher flip-flop density can be noticed for all the systems, dependent on the local lipid composition and often located away from the protein transmembrane domains (with the exception of COX1) (Figure 8 Figure Supplement 1). In contrast, flip-in events are concentrated in close proximity of the proteins and are not uniformly distributed but confined at distinct locations (Figure 8 Figure Supplement 1).

**Figure 7.**
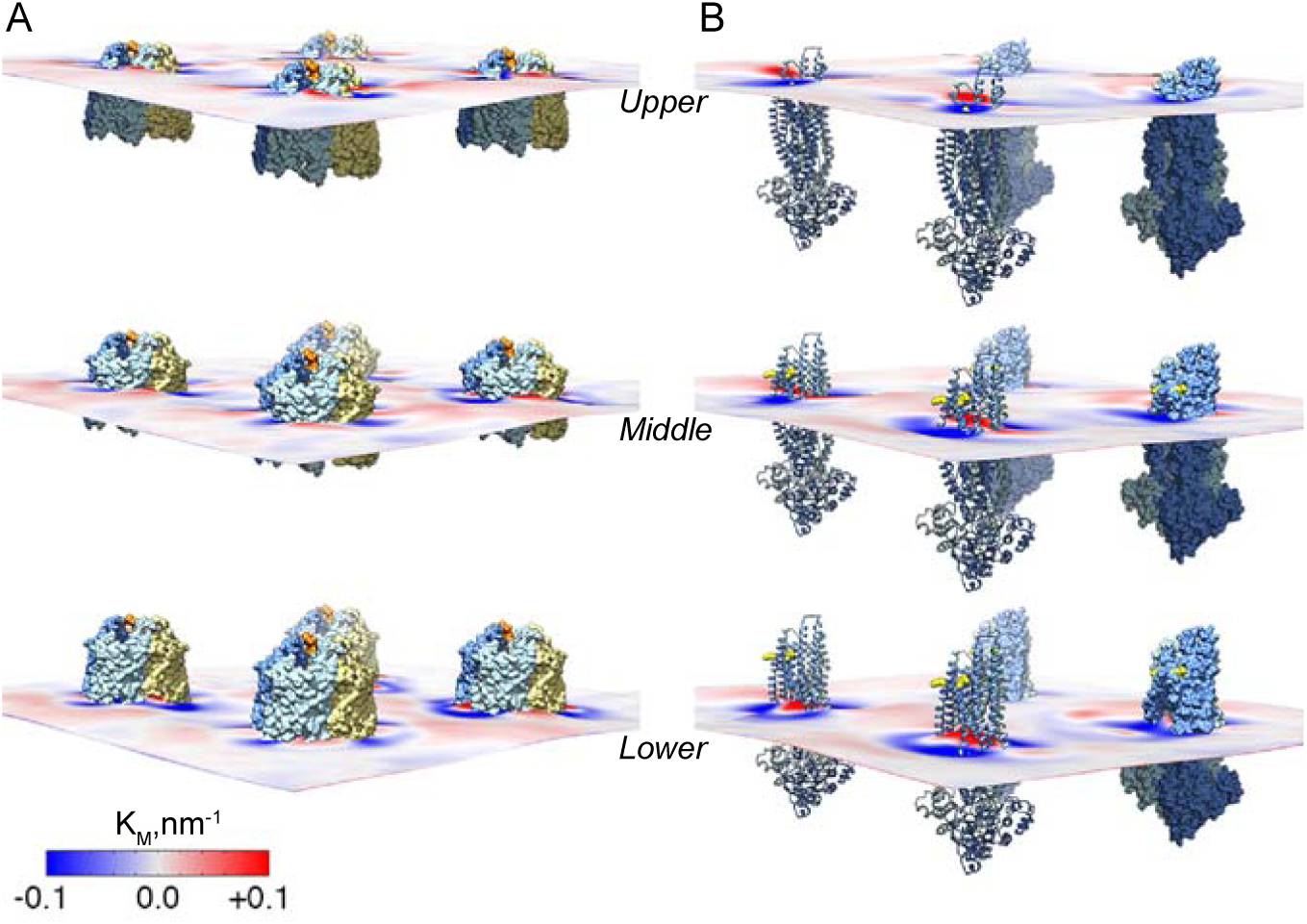
AQP1 and P-gp K_M_ profile. 3D representation of the four AQP1 (A) and P-gp (B) molecules and the 2D K_M_ maps for the upper, middle, and lower surface. Atomistic structures of AQP1 and P-gp were superimposed to the four corresponding CG protein molecules at 30 μs. In each AQP1 structure (A), the monomers are shown as orange, blue, lightblue and light-yellow surfaces. The two halves of each P-gp molecule (B) are shown in blueand light-blue cartoons for the molecules in the foreground, and surfaces for the protein in the background. Arg206 and Lys209 are represented in yellow spheres.

**Figure 8.**
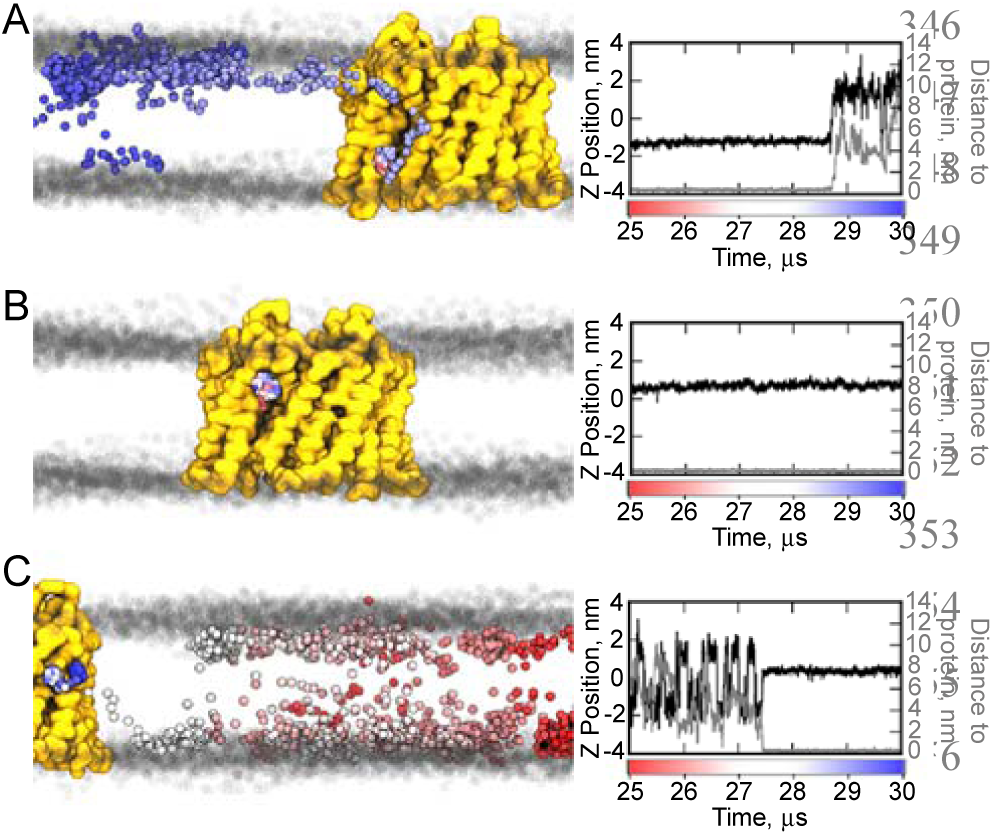
Cholesterol dynamics in the AQP1 system. (A-C) Representative examples of how cholesterol flip-flop might be affected by the presence of AQP1. In the left panels, for a given cholesterol molecule, the position of the ROH bead is shown every 2 ns from 25 to 30 μs, and colored from red to white to blue as a function of time. The AQP1 backbone is shown as a yellow surface, while the PO4 beads of the surrounding lipids are shown as grey spheres, taken every 1.0 μs during the 5 μs time. The graphs on the right show, as a function of time, the position of the corresponding ROH bead with respect to the middle of the bilayer (black line), and the minimum distance to the protein (grey line). The middle of the bilayer is set at 0 nm.

## Discussion

The function and mechanism of membrane proteins are modulated by lipid-protein interactions and dependent on membrane composition. AQPs, for example, are passive water channels whose function is critical to control cell volume and water balance. Such proteins are found associated with cholesterol-enriched domains,^43-51^ where cholesterol and lipid composition affect AQP-mediated water permeability. This has been shown, for example, for AQP4,^52^ normally expressed in brain astrocytes,^53-54^ characterized by a membrane with high cholesterol concentrations.^55-56^ Voltage-gated potassium channels (Kv) provide another example where protein function strictly depends on lipid composition and where the lipid environment dictates channel localization in the membrane.^57-67^ For several members of the Kv family, PIP lipids, and in particular PIP_2_, have been shown to modulate the kinetics of gating.^68^ In this complex lipid-protein interplay, beyond specific and non-specific lipid-protein interactions, geometrical properties of the membranes such as thickness and curvature are major players in regulating protein behavior.^37,69^

MD simulations have been used extensively to study how lipids, and in particular specific lipid-protein interactions, might regulate protein function, and recent advances in highperformance computing have allowed for a higher degree of complexity in the systems to simulate.^22-25, 29-34^ However, current limitations in this field still concern the tradeoff between complexity, timescale and statistical convergence of the results. Our approach considers four molecules of the same protein in a simulation system including a complex plasma membrane mixture, to account for reproducibility of the lipid distribution around a given protein type. The system size, although smaller compared to recently published studies,^30, 32^ allows for longer simulations, and was applied to explore the lipid organization for ten membrane protein types. We have tested different simulation setups and length, and obtained similar profiles in lipid enrichment near the proteins (Supplementary File 2, Table 1 Table Supplements 1 and 2).

Since the simulations are based on a complex and realistic plasma lipid mixture, with an asymmetric composition for the upper and lower leaflet, they contain a wealth of data about tendencies of different lipids to interact with different proteins, and in many cases different areas of a particular protein. A striking characteristic of the lipid composition maps (**Error! Reference source not found.** and Figure 3 Figure Supplements 1 and 2) is that there are some general features, but, overall, the distributions around each protein, and near different parts of each protein, are distinct, providing a unique environment or “lipid fingerprint” for each protein. Experimentally, many proteins are found associate with domains enriched in cholesterol and sphingolipids, and with higher content of FS lipids than the nearby domains.^70^

The presence of both poly-unsaturated and saturated lipid tails in contact with membrane proteins has been linked to membrane protein function regulation. Poly-unsaturated lipids, for example, are found in high concentrations in retinal membranes, where they modulate rhodopsin activity by stimulating the kinetics of the photocycle.^71-73^ Molecular dynamics simulations provided molecular details on the specific sites of lipid interactions, highlighting the different pattern of contacts of poly-unsaturated and saturated lipid tails.^74-75^ In addition, membrane protein sorting in regions enriched in unsaturated lipids has been observed in CG simulations of glycophorin A dimers, embedded in a red blood cell membrane model.^31^ In our study, the distribution of lipids classes around the proteins appears more complex. We observe the presence of regions of different size and composition, enriched in either FS or PU lipids, within few nm from the proteins and often with different distributions between upper and lower leaflet. As reported for the pure plasma membrane simulations,^35^ even in the presence of proteins we do not observe stable lipid domains. Experimental techniques such as electron spin resonance and differential scanning calorimetry revealed how membrane proteins play a significant role in sorting annular lipids and lipids at longer distances, as seen for example for the Ca^2+^-ATPase.^76-77^ Accordingly, in our simulations, lipids organized in stable regions enriched in FS or PU lipids (derived from averaging over a 5 μs window, **Error! Reference source not found.** and Figure 3 Figure Supplements 1 and 2, or over shorter time windows as in the case of AQP1, movie files), but such regions are strictly linked to the presence of the proteins, and anchored along the protein circumference. In the FS-enriched regions, the simulations reveal preponderant interactions between proteins and GM lipids. Glycolipid-protein interactions are involved in a number of cellular functions, as glycolipids-enriched domains participate to signal transduction and contribute to protein localization in the membrane.^78-79^ In our plasma membrane, glycolipid aggregation around the proteins is detected for all the systems (Figure 2 Figure Supplement 1, and Supplementary File 2, Table 1 Table Supplements 1 and 2), and although the magnitude of such aggregation varies from system to system, overall the glycolipid enrichment expands at least up to 2 nm from the proteins (Supplementary File 2, Table 1 Table Supplements 1 and 2). There is, however, limited experimental data available on specific glycolipid-protein interactions that could be used to validate the results. Protein function modulation induced by glycolipids interaction has been mainly described for receptors involved in signal transduction, including EGFR.^80^ However, previous simulation studies also highlighted the tendency of glycolipids to form small aggregates, and/or to interact with membrane proteins, at an atomistic or CG level of detail.^32, 34-35, 81-83^ While some of these studies used simplified membrane mixtures to study glycolipidprotein interactions, here we show the ability of CG simulations to retrieve such interactions in the context of a more complex plasma membrane mixture. These lipid-sorting events appear linked to the presence of membrane proteins, and may be in line with the lateral compartmentalization of the membrane, i.e. the glycolipid-enriched lipid raft hypothesis for protein localization and recruitment.^84^ However, we did not observe large-scale lipid sorting phenomena in our simulations, as glycolipid segregation lasting over tens of μs occurred only in close proximity of the proteins. In the lower leaflet, the membrane components that behave most similarly to the GM lipids of the upper leaflet are PIP lipids. Indeed, PIP lipids form small clusters in lipid bilayers and interact or bind with membrane proteins in many simulation studies.^30, 32, 34-35^ Here, common to most of the systems is a clear PIP lipids enrichment, which persists over few lipids shells around the proteins (Supplementary File 2, Table 1 - Table Supplements 1 and 2). Direct interaction between PIP lipids and membrane proteins has been shown for a number of channels and receptors, including EGFR and DAT, which are among the systems we simulated.^85-87^ However, given the variety of roles of this lipid type in the plasma membrane, from peripheral proteins localization, signaling, membrane trafficking and membrane protein function regulation,^88-89^ it is not surprising that the simulations detect interactions between PIP lipids and many other membrane proteins, thus providing some new details on possible specific lipid-protein interactions to investigate further.

The analysis of geometric properties of the bilayer, such as thickness and curvature, is relevant for protein function. The activity of a number of membrane proteins, including the Na,KATPase pump, potassium channels and others,^3, 90-92^ is tightly linked to hydrophobic mismatch between proteins and lipids. Values of hydrophobic thickness for membrane proteins vary significantly, from ca. 21 to ca. 44 Å.^93-94^ As a consequence, when various lipid species are present, the hydrophobic mismatch between lipids and proteins acts as one of the driving forces inducing depletion or enrichment of certain lipids. We observe, for example, proteins associated with regions of thinner membrane than others, as in the case of Kv1.2, one of the proteins with the smallest hydrophobic belt among those considered in this study, with a calculated hydrophobic thickness of ca. 25 Å.^95^ For Kv1.2 the thinning of the membrane is highly homogeneous around the entire tetramer, and extends across several lipid shells (**Error! Reference source not found.**). However, many other proteins are associated with both regions of increased and decreased thickness, as a consequence of a non-uniform protein hydrophobic belt, and the type of lipids associated with it. These regions of increased or decreased thickness span larger distances from the proteins (**Error! Reference source not found.** and Figure 5 - Figure Supplement 1), while previous MD studies have reported very homogenous thickness profiles when simulating proteins in model membranes of one or few lipid types.^96-99^ This would suggest that proteins facilitate the population of specific lipids in their neighborhood, which could then form sites from where larger lipid islands of similar type may form.

The protein-lipid interplay is the key factor in determining the shape of the membrane. The intrinsic flexibility of the lipids, and in particular the size of their head group can generate different spontaneous curvatures. Proteins, on the other hand, can bend membranes through a variety of mechanisms ^100^. Overall, the complex undulating profiles that we observe in our simulations (Figures 6-7 and Figure 6 - Figure Supplements 1 and 2) are the results of structural properties of the proteins, their shape, their depth of insertion, and the asymmetric distribution of lipids in the membrane, along with the clustering of certain lipid types (e.g. GM and PIP lipids). Membrane curvature is also a possible mechanism for communication between membrane proteins,^101^ and membrane protein oligomerization and redistribution.^69, 102-104^ For example, dimers of F_1_,F_0_-ATP synthase have been localized in the highly curved regions of mitochondrial membranes.^105^ Isolated dimers induce local deformations (curvature) spanning ca. 20 nm, which in turn drive the side-by-side organization of other dimers.^106^ Although the present study does not focus on protein oligomerization, we observe that membrane deformation in terms of curvature spans large distances, often connecting multiple protein copies, which in our systems are placed at ca. 20 nm distance. Considering the length-scale of membrane modification and its directionality, this study suggests the potential for collective effects/cooperative behavior in reshaping the membrane. Indeed, the formation of protein clusters has been shown to occur even in the absence of direct protein-protein interactions, simply driven by a certain degree of membrane curvature.^101^

## Conclusions

Combined, our simulations characterize the lipid fingerprint of each of the ten membrane proteins taken into account in this study. The lipid raft hypothesis outlines many membrane protein types as embedded in microdomains enriched in glycolipids, cholesterol and sphingolipids. However, the molecular picture resulting from our simulations describes a more complex and fragmented lipid environment, where regions enriched in different lipid classes coexist and rearrange around a given protein, and where long-lasting lipid segregation is mainly driven by direct interactions with the proteins. While general patterns can be observed, the molecular detail of this lipid environment is unique for each protein.

## Material and Methods

### Systems setup

We embedded ten different proteins in a previously characterized model plasma membrane.^35^ The proteins were: aquaporin 1 (AQP1),^107^ prostaglandin H2 synthase (COX1),^108^ dopamine transporter (DAT),^109^ epidermal growth factor (EGFR),^110^ AMPA-sensitive glutamate receptor (GluA2),^111^ glucose transporter (GLUT1),^112^ voltage-dependent Shaker potassium channel 1.2 (Kv1.2),^113^ sodium, potassium pump (Na,K-ATPase),^114^ δ-opioid receptor (δ-OPR),^115^ and P-glycoprotein (P-gp).^116^

Each protein structure, after removal of all the non-protein molecules, was converted in a CG model using the martinize protocol as described on Martini website (http://www.cgmartini.nl/), choosing the option of applying an elastic network on atom pairs within a 0.9 nm cut-off. One elastic network was applied when multiple chains were present, with the exception of AQP1, for which separate elastic networks were applied, one for each monomer of the tetramer. In the case of P-gp, the distance cut-off for the elastic network was increased to 1.0 nm, in order to include few elastic bonds between the two cytosolic domains. The initial simulation setup for GluA2 did not include the elastic network, which was added after 38 μs of simulation time, for additional 10 μs. DAT and GLUT1 were simulated with the presence of position restraints on the PO4 beads of selected phospholipids (POPC and PIPC in the upper leaflet), as in ^35^.

For each protein, the transmembrane region was identified using the corresponding entry of the OPM database.^95^ Four copies of each CG protein were placed in a simulation box of ca. 42x42 nm in x and y, and lipids, in a composition corresponding to the plasma membrane model developed by Ingólfsson and colleagues,^35^ were added using *insane*,^36^ for a total of ca. 6000 lipid molecules in each system. The following lipid classes were included: Cholesterol (CHOL), in both leaflet; charged lipids phosphatidylserine (PS), phosphatidic acid (PA), phosphatidylinositol (PI), and the PI-phosphate, -bisphosphate, and -trisphosphate (PIPs) were placed in the inner leaflet, and ganglioside (GM) in the outer leaflet. The zwitterionic phosphatidylcholine (PC), phosphatidylethanolamine (PE), and sphingomyelin (SM) lipids were placed in both leaflets, with PC and SM primarily in the outer leaflet and PE in the inner leaflet. Ceramide (CER), diacylglycerol (DAG), and lysophosphatidylcholine (LPC) lipids were also included, with all the LPC in the inner leaflet and CER, and DAG primarily in the outer leaflet.^35^ The exact lipid composition of each system is given in Supplementary File 1. Water molecules, counterions and 150 mM NaCl were also added.

### Simulation setup

Simulations were performed using the GROMACS simulation package version 4.6.x,^117^ with the standard Martini v2.2 simulation settings.^118^ After initial energy minimization with position restraints applied on the protein beads (using a force constant of 1000 kJ/mol nm^2^), short equilibrium runs were performed first with the position restraints applied to all the protein beads, and then to the backbone beads. All simulations were performed with a 20fs time step, a temperature of 310 K set using a velocity-rescaling thermostat^119^ with a time constant for coupling of 1 ps (2 ps for equilibrium runs). A semi-isotropic pressure of 1 bar maintained with the Berendsen barostat,^120^ with a compressibility of 3·10^-4^ bar^-1^, and a relaxation time constant of 5 ps. Production runs were performed in the presence of position restraints applied to the backbone beads, with a force constant of 1 kJ/mol nm^2^.

Following the systems and simulation setup described above and pictured in Figure 2, as an indication of the equilibration time of the number of specific lipids around a particular protein, the number of PC, GM and PIP lipids in contact with the proteins, as a function of time, was approximated by the number of PO4 (for PC), GM1 (for GM lipids) and CP (for PIP lipids) beads found within a 0.7 nm cut-off from the protein, similar to refs ^121-122^. The PC lipids were chosen as they are the most abundant phospholipid in the upper lipids, while GM and PIP lipids were selected as both tend to aggregate near the proteins (Figure 2 - Figure Supplement 1-3). The calculation was performed using the g_select tool implemented in GROMACS.^117^ The equilibration time is of the order of tens of microseconds. This information was used as a proxy for determining overall equilibration and is accurate for the larger groups of lipids, while rare lipids that are present in only a few copies do not yield very accurate distributions. Based on this, all the systems have been simulated for 30 μs. All the analyses, unless otherwise specified, were performed on the last 5 μs of each simulation system.

Additional control simulations were performed in order to test the effects of simulation length, lipid composition, and water model on the results of lipid composition near the proteins (Table 1 - Table Supplement 1). The AQP1 and Kv1.2 systems were extended to for 50 μs (Setup 1, Table 1 - Table Supplement 1); the AQP1 system was also simulated up to 50 μs (i) with position restraints on the headgroup of selected lipids and no restraints on the backbone beads (Setup 2, Table 1 Table Supplement 1); (ii) as Setup 2 but with no glycolipids in the mixture (Setup 3, Table 1 - Table Supplement 1); (iii) as Setup 2 but with polarizable water model^123^ (Setup 4, Table 1 - Table Supplement 1). Finally, the Na,K-ATPase system was simulated for an additional 20 μs after the initial 30 μs and after removing the glycolipids from the plasma mixture (Setup 5, Table 1 - Table Supplement 1).

### Analyses

#### Lipid Composition, Thickness, and Curvature

Thickness and curvature were calculated based on a method that uses three interpolated grid-surfaces (upper, middle, and lower). Surface averages are calculated for the last 5 μs on 30 μs long simulations, with a total of 2500 frames obtained by saving configurations every 2 ns. The three surfaces are defined using different lipid beads: PO4 and GM1 beads for the upper surface (plane); the last bead of each lipid tail for the middle surface; the PO4 beads for the lower surface. For the definition of these surfaces lipid species that do flip-flop during the simulations (CHOL, DAG, and CER lipids ^35^) were not taken into account. The choice of the GM1 bead for glycolipids was made with the help of small reference simulations (data not shown) consisting of binary mixtures of lipids (DPSM/DPG1) with equivalent acyl-chains and only differing in the headgroups. The GM4/GM1 beads of DPG1 have density peaks at positions equivalent to the DPSM-PO4 counterpart.

The method (to be published) has been implemented in C language, and has been derived from the numerical scheme described on a previous work that used MATLAB scripts,^124^ and where gradients of surfaces are defined by interpolation on squared grids with a previous averaging on molecular coordinates carried out by a Gaussian filter, used to eliminate noise and generate smooth surfaces. The grid spacing used was 0.3 nm, with a Gaussian filter that averages data for a maximum of 6 cells radii for every point on the grids.

Leaflet thickness was calculated via middle surface to upper/lower surface distance for every point in the grids. The overall thickness was likewise calculated as distance between the upper and lower surfaces.

The same surfaces defined for thickness calculation were used for the curvature analysis. The estimated spontaneous curvature of a grid patch is equivalent to the average curvature of the lipids in the upper surface, minus the average spontaneous curvature of the lipids in the lower surface, taking the lipid local normals to the membrane as a reference. Consequently, the curvature for the lower leaflet would have to be multiplied by minus one in order to find the correlation between membrane curvature and spontaneous curvature only for the lower leaflet.

Thickness values are given in nm, while mean and Gaussian curvatures are expressed on inverse distance units (nm^-1^ and nm^-2^, respectively).

Lipid composition is calculated by averaging the occupancy of cells for the entire set of 2500 frames, with units of lipid-tails per nm^2^. These values are then changed into density units of mass per unit of area by including lipid masses. The first tails defining the beginning of the acyl-chain on all lipids were used as criteria to decide the occupancy on every frame of the set of lipids selected. As in ^35^, four classes were analyzed, namely fully-saturated (FS), poly-unsaturated (PU), cholesterol (CHOL) and Others, with the last group defining lipids not present in the first three groups. The PU lipid class consists of DAPC, DUPE, DAPE, DAPS, DUPS, APC, UPC lipids (lipids where both the tails have more than two “D” type beads), while the FS class includes SM lipids (DPSM, DBSM, DXSM), glycolipids (DPG1, DXG1, DPG3, DXG3), ceramides (DPCE, DXCE), and LPC lipids (PPC).

For each class, the lipid composition was first calculated in terms of lipid density, and then changed into enrichment levels (*z_new_*) with respect to the average of the set (*z_ave_*). The new score, in percentage units, is defined by:

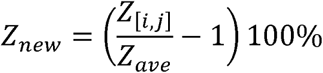

where the indices [*i*,*j*] correspond to every point in the grid to be reweighted. The new score has the particularity to be positive for *z_i,j_* values larger than *z_ave_*, and negative for values smaller than *z_ave_*. The 100% factor simply expresses the score as percentage units, indicating enrichment/depletion with respect to a homogeneous mixture with *z_ave_* score.

#### First shell lipid composition and depletion-enrichment index

The first shell lipid composition was calculated within 0.7 nm from the proteins. The depletion-enrichment (D-E) index of a given lipid type was calculated for three distance cut-offs from the protein, at 0.7, 1.4 and 2.1 nm. For a generic lipid type *L*, we first defined the ration of lipid *L* within a given cut-off *x* (namely *Ratio*(*L*)*_x_*), and the ratio of the lipid *L* with respect to bulk (namely *Ratio*(*L*)*_bulk_*) as follow:

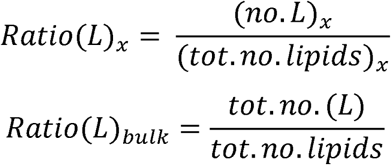

We used *Ratio(L)_x_* to calculate the fraction of lipid headgroup types (PC, PE, PS, PA, DAG, LPC, SM, CER, PI, PIPs, GM) present within 0.7 nm from the protein, during the last 5 μs of each simulation. For a given simulation system, which consists of four copies of the same protein, the lipid shell composition was calculated for each protein, and then averaged over the four protein copies.

The enrichment of the lipid *L* for a given cut-off *x* is then calculated from the following ratio:

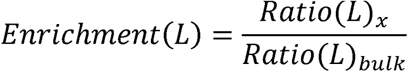

Selected beads were used to calculate the number of lipids within a cut-off *x* from any bead of the protein: the ROH bead was chosen for cholesterol, while GL1 or AM1 beads were used for all the other lipid types.

For all the systems, the enrichment was calculated for the last 5 μs for each individual lipid type for the upper and lower leaflet separately. For cholesterol, DAGs (PODG, PIDG, PADG, PUDG) and CERs (DPCE, DXCE, PNCE, XNCE), the analysis was performed by combining the two leaflets together, due to the possible flip-flop of these lipid species.

The enrichment was also calculated for groups of lipids categorized based on their headgroups (PC, PE, PS, PA, DAG, LPC, SM, CER, PI, PIPs, GM, GM1, and GM3) or tails (fully saturated lipids, poly unsaturated lipids and others). Here, a lipid is considered polyunsaturated if both the tails have more than two "D" type beads (DAPC, DAPE, DUPE, DAPS, DUPS, APC, UPC). In this case, the enrichment was calculated by combining the two leaflets together.

For a given simulation system, which consists of four copies of the same protein, the enrichment was calculated for each protein copy. The final values shown in Tables 1, and S1-3 correspond to the average values obtained from the enrichment values of the four protein copies. Standard deviations are also calculated.

#### Cholesterol dynamics

Cholesterol flip-flop and flip-in rates were calculated with a custom Python script that uses the MDAnalysis^125^ and NumPy^126^ packages. We used the PO4 beads of all the lipids in the upper and lower membrane leaflets to define those leaflets, and we considered a cholesterol molecule present in the upper/lower leaflet if its ROH bead was within 1.2 nm the respective PO4 bead group. Apart from cholesterol, CER and DAG lipids, the other lipid species in our simulation setup do not flip-flop at the simulation time scales ^35^, therefore, the predefined PO4 bead groups, and hence the definition of upper and lower leaflet for this analysis, do not change through the simulation.

To characterize cholesterol dynamics, we define flip-in and flip-flop events. A flip-in event is defined when a cholesterol ROH bead that used to be in either the upper or lower leaflet transitions into the bilayer middle (more than 1.2 nm away from the PO4 bead groups of either leaflets). A flip-flop event is defined when a cholesterol ROH bead that used to be in either the upper or lower leaflet transitions to the opposing leaflet. Both flip-flop and flip-in events were calculated from the last 5 μs of each simulation, using a trajectory with a frame rate of 2 ns. Flip-flop and flip-in rates per cholesterol were calculated as a function of the distance from the protein transmembrane domains, binned from 0 to 6 nm, with a bin widths of 0.5 nm. All events further than 6 nm from each of the four protein copies were classified as bulk. Additionally, the spatial distribution of flip-flop and flip-in in the membrane plain across a given simulation system is shown using x,y 2D density maps of the flip events. We calculated the number of events from 0 to 42 nm (0 to 36 nm for COX1), using a 2 nm bin width, and normalized by the number of cholesterol molecules in each bin.

## SUPPLEMENTARY FILES

Supplementary files include:

(a) Supplementary File 1 (Supplementary_File_1.xlsx), with:

Detailed composition of upper and lower leaflet for each system;

(b) Supplementary File 2 (Supplementary_File_2.xlsx), with:

Table 1 Table Supplements 1 and 2;

Table 2 Table Supplement 1;

(c) Movies S1-S4.

Movie S1: Enrichment-Depletion analysis movie for the PU, FS, CHOL, and Others lipid classes domains. This movie was obtained by averaging over 200 ns.

Movie S2: Enrichment-Depletion analysis movie for the PU, FS, CHOL, and Others lipid classes domains. This movie was obtained by averaging over 2000 ns.

Movie S3: Total thickness, upper and lower leaflet thickness movie obtained by averaging over 200 ns.

Movie S4: K_M_ and K_G_ for upper, middle and lower surfaces. This movie was obtained by averaging over 200 ns.

## AUTHOR INFORMATION

### Author Contributions

E. M.-V. wrote the tools to analyze lipid composition, thickness and curvature. H. I. I. and M. N. M. wrote the tools for the depletion-enrichment and cholesterol dynamics analysis. R.-X. G. carried out the depletion-enrichment analysis and simulated the Na,K-ATPase system. I. S., A. M., C. D., B. S. carried out the simulations of AQP1, DAT, GLUT1, Kv1.2 (I. S.); EGFR (A. M.); δ-OPR (C. D.); COX1 and GluA2 (B. S.). V. C. carried out the simulations of the P-gp system and analyzed the results from all the systems. G. S., T. A. W. and K. D. M. contributed with helpful discussions throughout the project. D.P.T. and S.J.M. designed the project. V. C., E. M.-V, D. P. T and S. J. M. wrote the manuscript, with contributions from H. I. I.

## ABBREVIATIONS

AQP1: aquaporin 1
COX1: prostaglandin H2 synthase
DAT: dopamine transporter
EGFR: epidermal growth factor
GluA2: AMPA-sensitive glutamate receptor
GLUT1: glucose transporter
Kv1.2: voltage-dependent Shaker potassium channel 1.2
Na,K-ATPase: sodium, potassium pump
δ-OPR: δ-opioid receptor
P-gp: P-glycoprotein (P-gp)
CHOL: cholesterol
PC: phosphatidylcholine lipids
PE: phosphatidylethanolamine lipids
SM: sphingomyelin lipids
PS: phosphatidylserine lipids
PA: phosphatidic acid lipids
PI: phosphati-dylinositol lipids
PIP: PI-phosphate, -bisphosphate, and -trisphosphate lipids
GM: ganglioside lipids
CER: ceramide
DAG: diacylglycerol lipids
LPC: lysophosphatidylcholine lipids
CG: coarse-grained
MD: molecular dynamics

